# miR-1/206 down-regulates splicing factor Srsf9 to promote myogenesis

**DOI:** 10.1101/691162

**Authors:** Kristen K. Bjorkman, Massimo Buvoli, Emily K. Pugach, Michael M. Polmear, Leslie A. Leinwand

## Abstract

**Background:** Myogenesis is driven by specific changes in the transcriptome that occur during the different stages of muscle differentiation. In addition to controlled transcriptional transitions, several other post-transcriptional mechanisms direct muscle differentiation. Both alternative splicing and miRNA activity regulate gene expression and production of specialized protein isoforms. Importantly, disruption of either process often results in severe phenotypes as reported for several muscle diseases. Thus, broadening our understanding of the post-transcriptional pathways that operate in muscles will lay the foundation for future therapeutic interventions.

**Methods:** We employed bioinformatics analysis in concert with the well-established C2C12 cell system for predicting and validating novel miR-1 and miR-206 targets engaged in muscle differentiation. We used reporter gene assays to test direct miRNA targeting and studied C2C12 cells stably expressing one of the cDNA candidates fused to a heterologous, miRNA-resistant 3’ UTR. We monitored effects on differentiation by measuring fusion index, myotube area, and myogenic gene expression during time course differentiation experiments.

**Results:** Gene ontology analysis revealed a strongly enriched set of putative miR-1 and miR-206 targets associated with RNA metabolism. Notably, the expression levels of several candidates decreased during C2C12 differentiation. We discovered that the splicing factor Srsf9 is a direct target of both miRNAs during myogenesis. Persistent Srsf9 expression during differentiation impaired myotube formation and blunted induction of the early pro-differentiation factor myogenin as well as the late differentiation marker sarcomeric myosin, Myh8.

**Conclusions:** Our data uncover novel miR-1 and miR-206 cellular targets and establish a functional link between the splicing factor Srsf9 and myoblast differentiation. The finding that miRNA-mediated clearance of Srsf9 is a key myogenic event illustrates the coordinated and sophisticated interplay between the diverse components of the gene regulatory network.

## Background

A complex network of integrated transcriptional and post-transcriptional regulatory mechanisms control skeletal muscle gene expression. Overall, skeletal muscle displays one of the most tissue-specific splicing profiles, and changes in alternative splicing trigger proper temporal gene expression patterns during myogenesis [1–7]. Moreover, biochemical and biophysical properties of many components of the contractile machinery can be fine-tuned by selectively expressing specialized isoforms. Regulated splicing is orchestrated by the combinatorial interaction between *cis*-regulatory elements and *trans*-acting factors. This process efficiently creates functionally diverse proteins from a single gene; for instance, the fast troponin T primary transcript can be alternatively spliced into 64 functionally distinct isoforms [8]. The critical importance of alternative splicing in skeletal muscle is even more apparent when this process is misregulated, as occurs in a number of muscle disorders [1]. For instance, perturbations in alternative splicing ultimately result in an inability to transition from a fetal to an adult splicing pattern in myotonic dystrophy 1 [9,10].

Members of both hnRNP and SR protein families play important roles during muscle development [11–13]. SR proteins constitute a highly conserved group of splicing factors required for both constitutive and alternative splicing [14,15]. They are structurally characterized by the presence of an N-terminal RNA recognition motif (RRM) and a C-terminal arginine/serine-rich domain (RS domain), which interacts with other cellular factors during spliceosome assembly and splice site selection. A number of reports underscore the vital importance of certain SR protein *in vivo*. For example, global SR gene knockouts exhibit embryonic lethality [13,16–19], and Srsf10 (Srp38) knockout embryos die mid-to late-gestation of multiple defects affecting both cardiac and skeletal muscle [13,19]. Finally, while conditional cardiac Srsf1 knockout animals die postnatally from defective juvenile-to-adult heart remodeling [18], conditional cardiac knockouts of Srsf2 (SC35) and Srsf4 (Srp75) are viable but develop dilated and hypertrophic cardiomyopathies, respectively [20,21].

While alternative splicing modulates the expression of different gene isoforms, miRNAs can control their cellular levels by base-pairing with miRNA response elements (MREs) present in target mRNAs. Formation of this duplex can subsequently block translation or trigger degradation of the mRNA. Importantly, disrupted miRNA processing in muscle leads to a perinatal lethal phenotype characterized by muscle hypoplasia, abnormal myofiber organization, and increased cell death [22]. Several specific miRNAs are enriched in skeletal muscle, including miR-1 and miR-206, which share a common seed sequence. These miRNAs suppress myoblast proliferation and promote muscle differentiation in both animal and cell culture models [23–27]. Both miR-1 and miR-206 are minimally expressed in proliferating myoblasts, but ectopic expression or inhibition can force or prevent differentiation, respectively [24–27]. Notably, miR-1 may also act as a key myogenic fate determinant as forced expression in HeLa cells shifts their transcriptome to a typical muscle profile [28].

miRNAs also control myogenesis by modulating the cellular concentration of several splicing factors. For example, miR-133a regulates nPTB, miR-30-5p targets MBNL, and miR-222 regulates Rbm24 [12,29,30]. In addition, the RNA helicases Ddx5 and Ddx17, which cooperate with the splicing factors hnRNP H/F and the transcription factor MyoD to regulate transcription of myogenic genes in C2C12 mouse myoblasts, are direct miR-1 and miR-206 targets [31].

Through a combination of *in silico* predictions and correlated myogenesis functional analyses in C2C12 cells, we report herein that miR-1 and miR-206 targeting of the transcript encoding the SR protein Srsf9 is a significant myogenic event. The data presented highlight the cellular relationship between miRNA expression and levels of splicing factors that control the temporal production of specialized isoforms during muscle differentiation.

## Methods

### RNA isolation, cDNA synthesis, and qPCR

Total RNA was isolated with TRI Reagent (MRC TR 118) according to the manufacturer’s protocol. Tissue samples were homogenized directly into TRI reagent using a dispersion tool (IKA T10 Basic S1) while cultured cells were washed with PBS and scraped directly into TRI reagent. All qPCR was performed on a Bio-Rad CFX96 attached to a C1000 thermocycler. miRNA expression was assessed by TaqMan-based qPCR (ThermoFisher, Part Number 4427975; Assay IDs: sno202 = 001232, miR-206 = 000510, miR-1 = 002222). miRNAs were reverse transcribed with a TaqMan MicroRNA Reverse Transcription Kit (ThermoFisher, 4366596) and qPCR was performed with TaqMan Universal PCR Master Mix, No AmpErase UNG (ThermoFisher, 4324018), all according to the manufacturer’s instructions. Relative expression was analyzed with the ΔΔC_T_ method. mRNA expression was assessed with SYBR Green-based qPCR. All primer sequences are listed in Supplementary Table 1. Total RNA was reverse transcribed with random hexamer primers using the Superscript II Reverse Transcriptase kit (ThermoFisher, 18064-022). qPCR was performed with SYBR Green PCR Master Mix (ThermoFisher, 4312704) according to the manufacturer’s instructions. Relative expression was analyzed with the Pfaffl standard curve method.

### RNA sequencing

Total RNA from 3 independent growing myoblast cultures before differentiation (BD1, BD2, BD3) and 3 independent day 6 myotube cultures after differentiation (AD1, AD2, AD3) was oligo dT-selected and reverse transcribed with random hexamers, followed by second strand synthesis, adapter ligation, and 5’ end selection. Libraries were sequenced with an Illumina HiSEQ. 210 million paired end reads were aligned to the mm9 version of the mouse genome downloaded from UCSC Genome Browser. Supplementary Table 2 presents reads per million aligned. BAM files are available upon request.

### miR-1/206 target selection and gene ontology (GO) analysis

TargetScan 7.2 was queried to retrieve the set of predicted miR-1/206 mouse target genes [32]. This was crossed with a list of all mRNAs in the RNA-Seq dataset that decreased by 1.3-fold or more during differentiation, regardless of statistical significance. These permissive criteria were chosen to be as inclusive as possible for potential miR-1/206 targets in C2C12. This list was analyzed with DAVID (Database for Annotation, Visualization and Integrated Discovery version 6.8 [33–35]) to uncover enriched GO clusters of related gene sets. The functional clustering tool was used with default settings and the mouse genome as background. Clusters were ranked by descending Enrichment Score. Enrichment score is the –log_10_ of the geometric mean of the p-values of all individual terms in the cluster. A cutoff of 0.05 or below for the geometric mean of p-values, which corresponds to an Enrichment Score of greater than 1.3, was chosen.

### Cloning and mutagenesis

All primer sequences are listed in Supplementary Table 1. miRNA overexpression constructs were generated by cloning either the stem loop region (miR-450a-1 and miR-1) or a larger 1 kilobase region encompassing the stem loop (miR-206; the stem loop sequence alone was insufficient for proper processing and subsequent targeting; data not shown) into pcDNA3.1(-) through standard molecular cloning techniques. miR-450a-1 and miR-1 stem loops were assembled with primers through overlap extension PCR and cloned into the EcoRI/BamHI sites of the plasmid while the 1 kb miR-206 locus was PCR-amplified from mouse genomic DNA and inserted between the HindIII and XhoI sites of the plasmid. 3’ UTR reporter constructs were generated in psiCheck2, a dual luciferase plasmid where UTRs are cloned downstream of Renilla luciferase and firefly luciferase serves as an internal control. All sequences were cloned between the XhoI and NotI sites. Positive controls for miR-450a-1, miR-1, and miR-206 (2×450a-1, 2×1, and 2×206) were constructed by introducing 2 repeats (spaced by 2 A residues) of the mature miRNA antisense sequence downstream of Renilla luciferase using annealed complementary oligonucleotides. 3’ UTRs were PCR-amplified from mouse genomic DNA and inserted into the plasmid using standard molecular cloning techniques. The Srsf9 MRE was mutated by replacing the natural MRE with the reverse sequence through inverse PCR of the entire psiCheck2-Srsf9 wild-type 3’ UTR plasmid. iProof high fidelity DNA polymerase (Bio-Rad, 172-5302) was used according to the manufacturer’s instructions with phosphorylated primers. pEGFP-Srsf9 (a C-terminally GFP-tagged Srsf9 expression construct) was generated by amplifying Srsf9 cDNA from C2C12 myoblast cDNA and inserting it between the EcoRI and BamHI sites of pEGFP-N1 with standard molecular cloning techniques. A C-terminal GFP tag in a related vector, pEGFP-N3, has been shown to be tolerated by other SR proteins [36]. All clones were verified by Sanger sequencing.

### Cell culture, transfection, and stable cell line generation

C2C12 cells were grown as myoblasts in Growth Medium (GM): high glucose DMEM (Invitrogen 11960069) supplemented with 20% fetal bovine serum, 2 mM L-glutamine, 100 U/mL penicillin and 100 μg/mL streptomycin, and 1 mM sodium pyruvate. Myotubes were differentiated by changing media to Differentiation Medium (DM): high glucose DMEM supplemented with 5% adult horse serum, 2 mM L-glutamine, 100 U/mL penicillin and 100 μg/mL streptomycin, and 1 mM sodium pyruvate. When differentiating, DM was refreshed every day to prevent media acidification. For luciferase assays, cells were plated in triplicate in 6-well dishes at a density of 50,000 cells/well 24 hours before transfection. Cells were transfected 24 hours after plating using the transfection reagent *Trans*IT-LT1 (Mirus, MIR 2305) according to the manufacturer’s instructions. When ectopically expressing a miRNA along with a 3’ UTR-linked reporter gene, cells were harvested 24 hours post-transfection. When differentiating cells for endogenous induction of miRNAs, Day 0 timepoints were collected 24 hours post-transfection and differentiation was triggered at the same time for later timepoints. At harvest, cells were washed twice with phosphate-buffered saline (PBS) solution and stored in an ultralow freezer in order to process all timepoints together at the end of the experiment. For stable cell line generation, C2C12 cells were transfected with plasmids (either pcDNA3.1(-), pEGFP-N1, or pEGFP-N1-Srsf9) using *Trans*IT-LT1. 24 hours post-transfection, G418 antibiotic selection was initiated by adding GM supplemented with 250 μg/mL G418. Selection was continued until control untransfected cells all died and cells were split whenever necessary to keep density below 60% to prevent spontaneous differentiation. Cell lines were maintained as pools to minimize potential genomic locus insertional effects.

### Tissue collection

Animal work was reviewed by the University of Colorado Boulder Institutional Animal Care and Use Committee and approved under protocols 1002.07 and 1002.08. Wild-type C57Bl/6 mice were bred and housed at the University of Colorado Boulder under standard conditions. Adult animals were anesthetized by inhaled isoflurane and sacrificed by cervical dislocation followed by pneumothorax. Embryos were anesthetized on ice and sacrificed by decapitation. Whole hindlimbs were collected for embryonic timepoints and males and females were equally represented (sex was determined by PCR for the Sry gene using genomic DNA from the tail as template and primers against β-glucuronidase on chromosome 5 as positive control; see Supplementary Table 1 for primer sequences). Adult soleus was collected from 6 month-old male mice. Tissue samples were flash frozen in liquid nitrogen and stored in an ultralow freezer. Animal numbers for 15.5 dpc, 17.5 dpc, 19.5 dpc, and adult were 4, 6, 6, and 4, respectively.

### Luciferase assays

A Dual-Luciferase Reporter Assay System (Promega, E1960) was used according to the manufacturer’s instructions with a Turner Designs TD-20/20 luminometer. Briefly, frozen cells were lysed in 1X Passive Lysis Buffer and reporter gene activities were measured first in LARII reagent (firefly luciferase internal control) and second in Stop & Glo reagent (Renilla luciferase experimental reporter gene). Read times for both were 10 seconds. Renilla/firefly ratios were compared.

### Western blotting

Cell lysates were prepared in ice cold RIPA buffer (50 mM Tris pH 8.0, 1 mM EDTA, 0.5 mM EGTA, 1% Triton X-100, 0.5% deoxycholate, 0.1% SDS, 140 mM NaCl) with 1X cOmplete EDTA-free Protease Inhibitor Cocktail (Roche, 11873580001). Total protein concentration was measured with a BCA protein assay kit (Pierce, 23250) and 20 μg total protein was resolved by 10% PAGE and transferred to nitrocellulose membrane. Standard western blotting techniques were employed with all antibodies diluted in TBS/0.1% tween-20/4% nonfat dry milk. 1:2,000 α-GFP (Santa Cruz Biotechnology, sc-8334, rabbit polyclonal) and 1:5,000 α-Gapdh (Cell Signaling Technology, 2118S, rabbit monoclonal 14C10) were used as primary antibodies and horse radish peroxidase-linked goat anti-rabbit (Jackson Immunoresearch, 111-035-144) was used as secondary.

### Immunostaining and microscopy

Cells were seeded at a density of 100,000 cells per well in 6-well dishes on glass coverslips that had been silanized, gelatinized, and glutaraldehyde-crosslinked to promote adhesion of myotubes. Cells were harvested by washing once in 37°C PBS + 0.4% glucose and fixed 5’ at room temperature in 3% paraformaldehyde diluted in PBS. Paraformaldehyde was quenched with 50 mM ammonium chloride. Fixed cells were permeabilized 5’ at room temperature in PBS/0.1% triton X-100 (Permeabilization Solution (PS)) and blocked in Blocking Solution (BS: PS/1% BSA/1% normal goat serum) while rocking. Myosin heavy chain was probed with F59 as primary antibody (Developmental Studies Hybridoma Bank; custom production from hybridoma cells) and AlexaFluor568-linked goat anti-mouse IgG1 as secondary antibody (Invitrogen, A-21124) diluted 1:200. DNA was stained with 300 nM 4′,6-diamidino-2-phenylindole (DAPI; Sigma D9542). All imaging (for GFP, AlexaFluor568, and DAPI) was performed on a Nikon TE2000 inverted fluorescent microscope with a 20X objective connected to a Nikon DS-QiMc-U3 camera controlled through the NIS-Elements AR software version 4.00.03. Exposure times for a given channel were kept constant for all slides. Raw images were processed with the Image5D plugin in ImageJ using the same settings for a given channel across all images.

### Nuclear fusion index and myotube area calculation

Fusion indices were calculated from the percent of total nuclei residing in syncytia (syncytium defined as a cell with 2 or more nuclei). 8-10 non-overlapping 20X fields of view were analyzed and averaged for each cell line. The number of nuclei in syncytia was tabulated and divided by the total nuclei in each field. Myotube area per field of view was calculated as follows. Red (myosin) and blue (DNA) channels for a given field of view were individually opened in ImageJ. Each was inverted and thresholded (the same threshold for a given channel was applied to all images) to create a binary black and white mask. The Analyze Particles function was run on each myosin and DNA image and the area of the myosin image was divided by the area of the DNA image (to normalize for potential differences in cell density across cell lines and across different fields of view). A representative set of thresholded image masks is displayed in Supplementary Figure 1.

### Graphing and statistical analysis

All graphing was done with GraphPad Prism and statistical significance was assessed by one-way ANOVA with a Dunnett’s post-test to compare to the control condition or a Tukey’s post-test to compare all pairs as noted for multiple comparisons. Data are presented as means with error bars representing standard error of the mean. To assess potential differences in myogenin and perinatal myosin heavy chain expression over a differentiation time course, nonlinear regression analysis was used to fit quadratic curves to the data points for GFP control and Srsf9-GFP cell lines. Curve fits were compared to determine whether they were statistically different. A p-value cutoff of 0.05 was the minimum for significance. Other cutoffs are as noted in the figure legends.

## Results

### Splicing factors are enriched amongst predicted miR-1/206 targets and they decrease in expression during myotube formation

In order to identify functional categories of putative miR-1/206 targets in myogenesis, we took a combined *in silico* and *in vitro* cell culture model approach. We reasoned that miR-1/206 target genes would be expressed in myoblasts as they are poised to differentiate but that their concentrations would be reduced during the differentiation process by the increasing levels of miR-1/206. To this end, in both C2C12 myoblasts and day 6 myotubes we initially measured the amounts of miR-1 and miR-206 by qPCR as well as the global changes in their mRNA transcriptomes by RNA-Seq (Supplementary Table 2). These analyses showed that while miR-1 and miR-206 levels are very low in myoblasts, both miRNAs are robustly expressed in differentiated myotubes, which have both the molecular and morphological characteristics of mature muscle (Supplementary Figure 2, Panels A and B). This also confirms previous measurements of miR-1 and miR-206 in differentiating C2C12 cells [26]. Since miRNAs often exert mild effects on individual targets [37,38], we used the RNA-Seq dataset to assemble a list of genes down-regulated by 1.3-fold or more during differentiation. We then retrieved the set of 896 predicted miR-1/206 targets from TargetScan and found that the levels of 354 decreased during myogenesis (Supplementary Table 3).

In order to evaluate whether particular functional categories are enriched in this gene set, we employed the DAVID functional annotation clustering tool, which groups functional categories with similar gene sets from multiple different annotation sources to capture biological themes. Of the 354 candidate C2C12 miR-1/206 targets, 351 had associated DAVID IDs. We found 21 clusters (Supplementary Table 4) with an enrichment score greater than 1.30 (indicating an average p-value of below 0.05 for all terms in the cluster). While the top two clusters were related to DNA binding and transcriptional regulation and included known targets such as E2f5 [39], Pax3 [40], and Sox9 [41], both the third and fifth clusters included genes related to RNA metabolism. Since regulated splicing plays a central role in promoting myogenesis [4,6,7], we investigated whether miR-1/206 control muscle differentiation by specifically targeting RNA processing factors.

### mRNAs encoding SR protein family members are miR-1/206 targets during myogenesis

Our bioinformatics analysis predicted mRNAs encoding four of these proteins, Srsf1 (ASF/SF2), Srsf3 (Srp20), Srsf9 (Srp30c), and the SR-related protein Tra2b, as miR-1/206 targets. Since Srsf1 has an established critical role in heart development [18], we focused our attention on the remaining candidates. We assayed miRNA targeting in C2C12 myoblasts, which, as already noted, have extremely low levels of both miR-1 and miR-206. We co-transfected cells with constructs expressing a reporter Renilla luciferase gene fused to the mouse 3’ UTRs of our RNA candidates and either miR-1 or miR-206 overexpression plasmids. We determined the specificity of miR-1 or miR-206 by measuring the targeting activity of miR-450a-1, a non-myogenic miRNA which does not have overlapping predicted targets with miR-1/206. Luciferase quantification showed that this miRNA did not cross-react with the selected SR 3’ UTRs or with 3’ UTRs carrying tandem copies of the complete reverse complement of either miR-206 (2×206) or miR-1 (2×1) (data not shown). However, it efficiently down-regulated the positive control construct 2×450a-1, constructed similarly to 2×206 and 2×1 (Supplementary Figure 3). To determine the sensitivity of the assay, we next assessed the activity of miR-1/206 on the 2×206 and 2×1 constructs as well as on the 3’ UTR of Ccnd1, previously identified as a miR-206 target in C2C12 cells [42]. We also monitored promoter interference causing potential transcriptional repression of the reporter construct using a miRless construct, which expresses the Renilla luciferase RNA without miRNA target sites. Figure 1A shows the result of this analysis with each luciferase signal measured in the presence of miR-1 or miR-206 normalized to the same constructs co-transfected with the control miRNA, miR-450a-1 (see Materials and Methods). The data, presented as fold change versus miR-450a-1 control, show that miRless expression did not change appreciably in the presence of miR-1 or miR-206 and that miR-1 and miR-206 efficiently targeted 2×1 and 2×206 along with the positive control Ccnd1 (Figure 1A). Furthermore, both miR-1 and miR-206 also reduced luciferase activity by targeting the 3’ UTRs of Srsf9 and Tra2b fused to the reporter gene while they did not have any effect on the Srsf3 construct (Figure 1B). While statistically significant, the negative regulation of the Tra2b 3’ UTR was modest in magnitude. Thus, we focused our next set of experiments only on Srsf9 activity and its potential role in muscle differentiation. We substantiated the specific miR-1/206 targeting of the Srsf9 3’ UTR by MRE mutagenesis. When we reversed the orientation of the predicted MRE in the Srsf9 3’ UTR, which preserves positioning of any unrecognized flanking elements, this mutant construct (termed Srsf9 MRE Rev) restored reporter gene activity to levels measured in the presence of the control miR-450a-1 (Figure 1C). Taken together, these results establish that expression of the miR-1/206 family can directly modulate the level of Srsf9 in C2C12 myoblasts.

**Figure 1.**
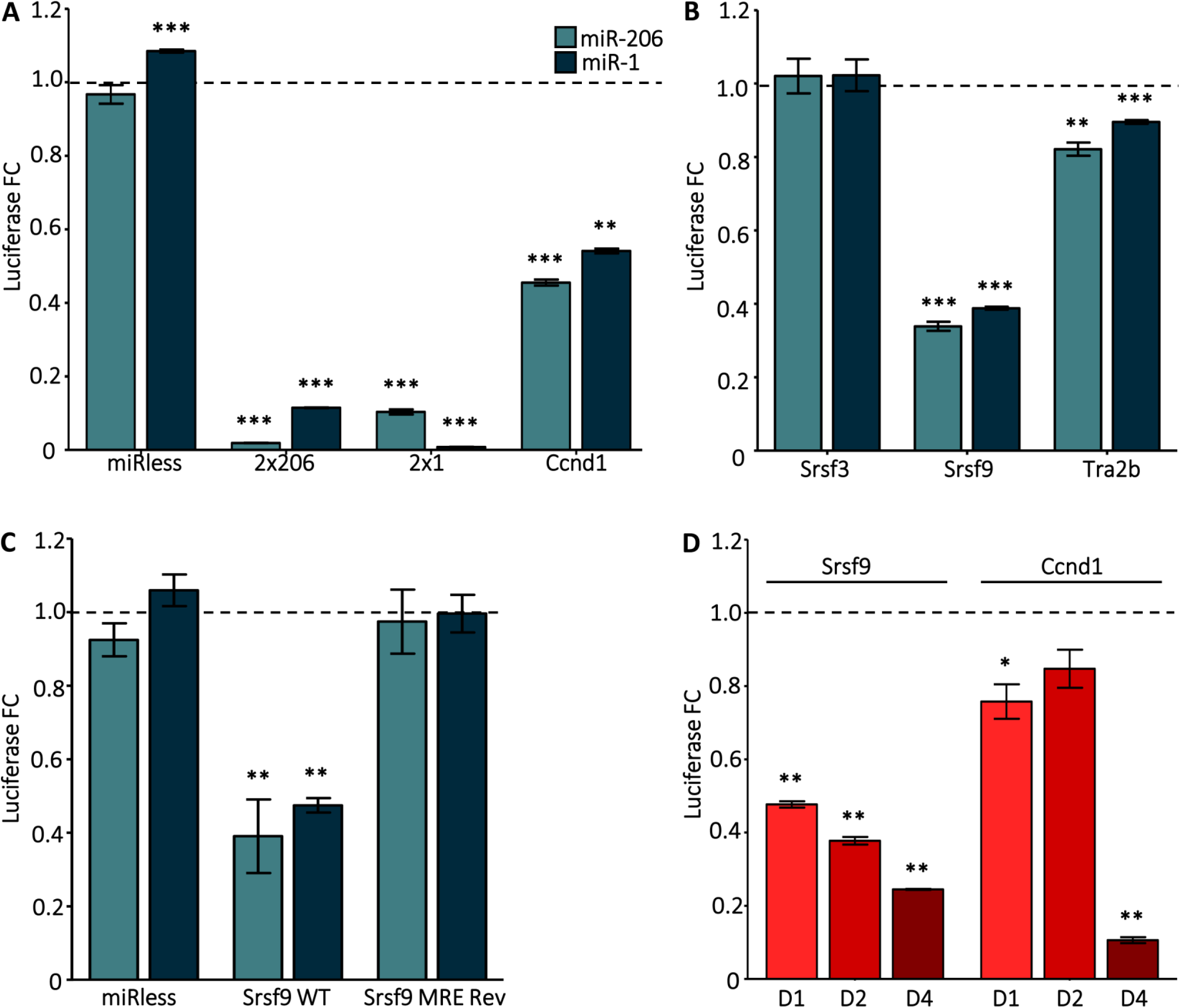
Srsf9 is a direct target of miR-1 and miR-206. **(A)** miR-206 and miR-1 target 3’ UTRs with cognate MREs in C2C12 myoblasts. Cells were transfected with reporter constructs and expression constructs for miR-206, miR-1, or the control miRNA, miR-450a-1. Renilla luciferase linked to two tandem copies of the reverse complement of miR-206 or miR-1 (2×206 and 2×1, respectively) or to the 3’ UTR of known miR-1/206 target Ccnd1 were positive controls. A 3’ UTR with no MREs in the empty reporter construct (miRless) was the negative control. Cells were harvested 24 hr after transfection. The graph presents fold changes (FC) in firefly-normalized Renilla luciferase activity for miR-206-or miR-1-transfected vs. miR-450a-1-transfected cells. The dashed line at y=1 represents miR-450a-1-transfected levels. ** = p ≤ 0.01; *** = p ≤ 0.001 vs miR-450a-1 **(B)** 3’ UTRs from Srsf9 and Tra2b but not Srsf3 are sensitive to miR-206 and miR-1 activity. C2C12 myoblasts were transfected with reporter constructs containing the Srsf3, Srsf9, or Tra2b 3’ UTRs along with miRNA expression constructs. Data were collected and analyzed as in A. **(C)** Srsf9 is a direct target of miR-1/206. The predicted MRE in the Srsf9 3’ UTR was reversed in the reporter construct to abolish miR-1/206 binding. C2C12 myoblasts were transfected and analyzed as in Panel A. **(D)** The Srsf9 3’ UTRs is sensitive to endogenous myogenic cues. 3’ UTR reporter constructs were transfected into C2C12 myoblasts which were then differentiated for 0, 1, 2, or 4 days. Firefly-normalized Renilla luciferase activity was measured. Fold changes for differentiating vs. Day 0 myoblasts were calculated and normalized to the miRless control. The dashed line at y=1 represents Day 0 levels. * = p ≤ 0.05; ** = p ≤ 0.01 vs D0 For all panels, N = 3 independent cultures.

To broaden the data obtained in myoblasts, we next surveyed the activity of the reporter construct during C2C12 differentiation which exposes the Srsf9 3’ UTR to physiological concentrations of both miR-1 and miR-206. Time course analysis showed an inversely proportional relationship between the decreased reporter luciferase activity and the concomitant increase in miR-1 and miR-206 expression occurring during the activation of the C2C12 differentiation program (Figure 1D; Supplementary Figure 2). Interestingly, we observed a similar pattern for the Ccnd1 reporter construct (Figure 1D). Consistently, the mRNA expression levels of Srsf9 and Ccnd1 measured six days after differentiation were significantly reduced (Figure 2A), which is also in accordance with the RNA-Seq data. We also queried a quantitative pSILAC mass spectrometry dataset that compiles protein level changes in HeLa cells 8 hr after miR-1 overexpression. The median fold change for Srsf9 peptides was −3.71, which supports direct miR-1 targeting [37,43].

**Figure 2.**
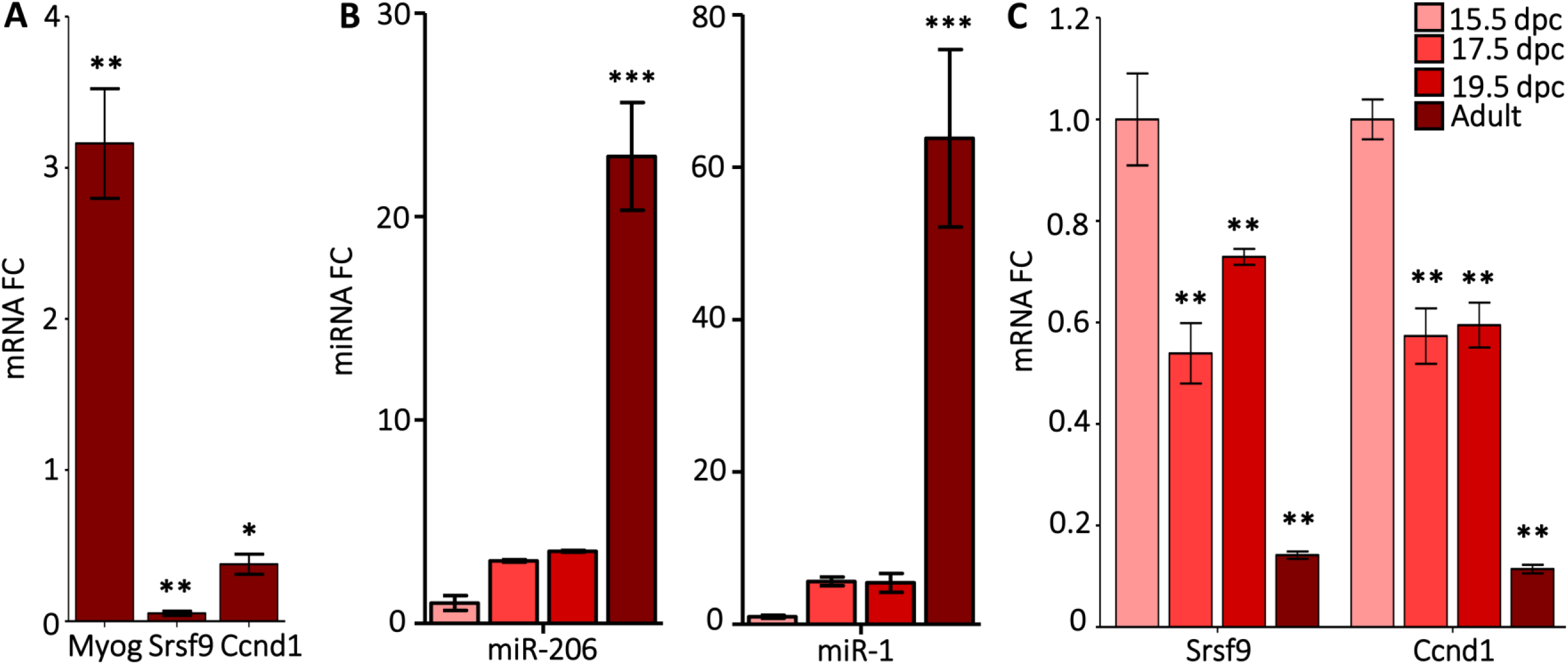
Srsf9 mRNA levels are inversely proportional to miR-1/206 levels. **(A)** Srsf9 mRNA levels decrease in differentiating C2C12 cells. Expression was assessed by qPCR in growing myoblasts (Day 0) and in Day 6 myotubes. Gapdh was the reference gene. Expression levels in the differentiating samples were normalized to myoblast levels (indicated by the dashed line). Myogenin (MyoG) was the positive control as its expression increases during differentiation. * = p ≤ 0.05; ** = p ≤ 0.01 vs. D0 N = 3 independent cultures for all genes and timepoints. **(B)** miR-206 and miR-1 levels increase during mouse hindlimb skeletal muscle development. Expression was assessed by qPCR and normalized to sno202 in whole hindlimb for developmental timepoints and in the soleus from 6 month-old male mice. Fold change relative to 15.5 dpc is presented. N for 15.5 dpc, 17.5 dpc, 19.5 dpc, and adult was 4, 6, 6, and 4, respectively. *** = p ≤ 0.001 vs. 15.5 dpc **(C)** Srsf9 and Ccnd1 mRNAs decrease during mouse hindlimb skeletal muscle development. Expression was assessed by qPCR with 18S rRNA as the reference gene. Expression levels were normalized to 15.5 dpc (indicated by the dashed line). * = p ≤ 0.05; ** = p ≤ 0.01 vs 15.5 dpc Ns were the same as in Panel B.

We extended this analysis to embryonic limb formation in the developing mouse. As shown in Figure 2B, we found that both miR-1 and miR-206 expression, which increased between 3- and 5-fold as embryogenesis progressed, peaked during the transition to mature adult muscle. In agreement with the inverse correlation observed in the C2C12 analysis, the mRNA levels of Srsf9 and Ccnd1 showed a modest decline in expression from embryonic day 15.5 to day 19.5 but a marked decrease from the embryonic to adult mouse transition (Figure 2C).

### The inability to down-regulate Srsf9 expression during myogenesis results in impaired differentiation

Finally, we investigated whether the down-regulation of Srsf9 expression plays an important role in controlling the sequential myogenic pathway. To this end, we generated C2C12 stable cell lines expressing a GFP-tagged version of Srsf9 fused to a heterologous 3’ UTR containing the SV40 polyA signal as well as GFP only or empty vector controls. After selection, we maintained pools of stable transfectants and measured transgene RNA and protein expression during differentiation. qPCR and western blot analysis carried out with GFP-specific primers and antibody revealed that both Srsf9-GFP and GFP were stably expressed in our cell pools (Supplementary Figure 4 A,B). More importantly, time course analysis showed that the presence of the heterologous 3’ UTR stabilized the level of Srsf9-GFP mRNA throughout cell differentiation (Supplementary Figure 4C). Moreover, we found that, as revealed by co-localization with DAPI stain, Srsf9-GFP was properly localized in the nucleus of the majority of the cells, while the GFP control showed diffuse fluorescence signal (Figure 3A). To determine potential functional repercussions caused by sustained Srsf9 expression, we first calculated the fusion index, a method frequently used to quantify the extent of C2C12 differentiation. Scoring of the number of nuclei residing in a fused syncytium (defined as a cell with 2 or more nuclei) at day 6 of differentiation showed that the average fusion index of cells expressing Srsf9-GFP was significantly lower than the ones expressing either GFP or the empty vector control (Figure 3B). Since Srsf9-GFP differentiated myotubes imaged at day 6 appeared to have reduced width compared to controls, we measured whether the myotube-positive area of each field of view was statistically smaller when normalized to DNA-positive area, a control for potential differences in cell density. After imaging the different cell pools stained with DAPI and a myosin heavy chain antibody (considered a terminal marker of muscle differentiation), we created cell masks by imposing equal intensity thresholds to the acquired images and then divided the myosin area by the DAPI area derived from the binary derivatives. As shown in Figure 3C, the stable expression of the Srsf9-GFP construct, which is refractory to miR-1/206 targeting activity, resulted in a significant reduction of myotube area, a clear indication of impaired differentiation. Supporting this observation, induction of the perinatal myosin isoform (Myh8), the most expressed myosin in mammalian skeletal muscle during the early perinatal period [44] and one of the most abundant myosins expressed in differentiated C2C12 (Supplementary Table 2), was significantly blunted in Srsf9-GFP cells (Figure 3D). This is indicated not only by decreased expression at individual timepoints but also by statistically different curves fit to each series when comparing GFP control and Srsf9-GFP cells. Finally, as the myogenesis program is fundamentally orchestrated at the transcriptional level, we measured induction of the muscle regulatory factor myogenin which is robustly expressed at early myogenic timepoints and then decreases as myotubes approach terminal differentiation. Although both the GFP controls and the Srsf9-GFP cells induced myogenin relative to D0 growing myoblast expression levels, this induction was much weaker in the Srsf9-GFP cells (Figure 3E). Since the series could still be fit to a quadratic curve, with myogenin expression in Srsf9-GFP cells peaking mid-time course and then declining at day 6, we believe the data do not support a simple delay of differentiation. Taken together, these data strongly suggest that proper progression throughout the steps of muscle differentiation requires temporal Srsf9 down-regulation.

**Figure 3.**
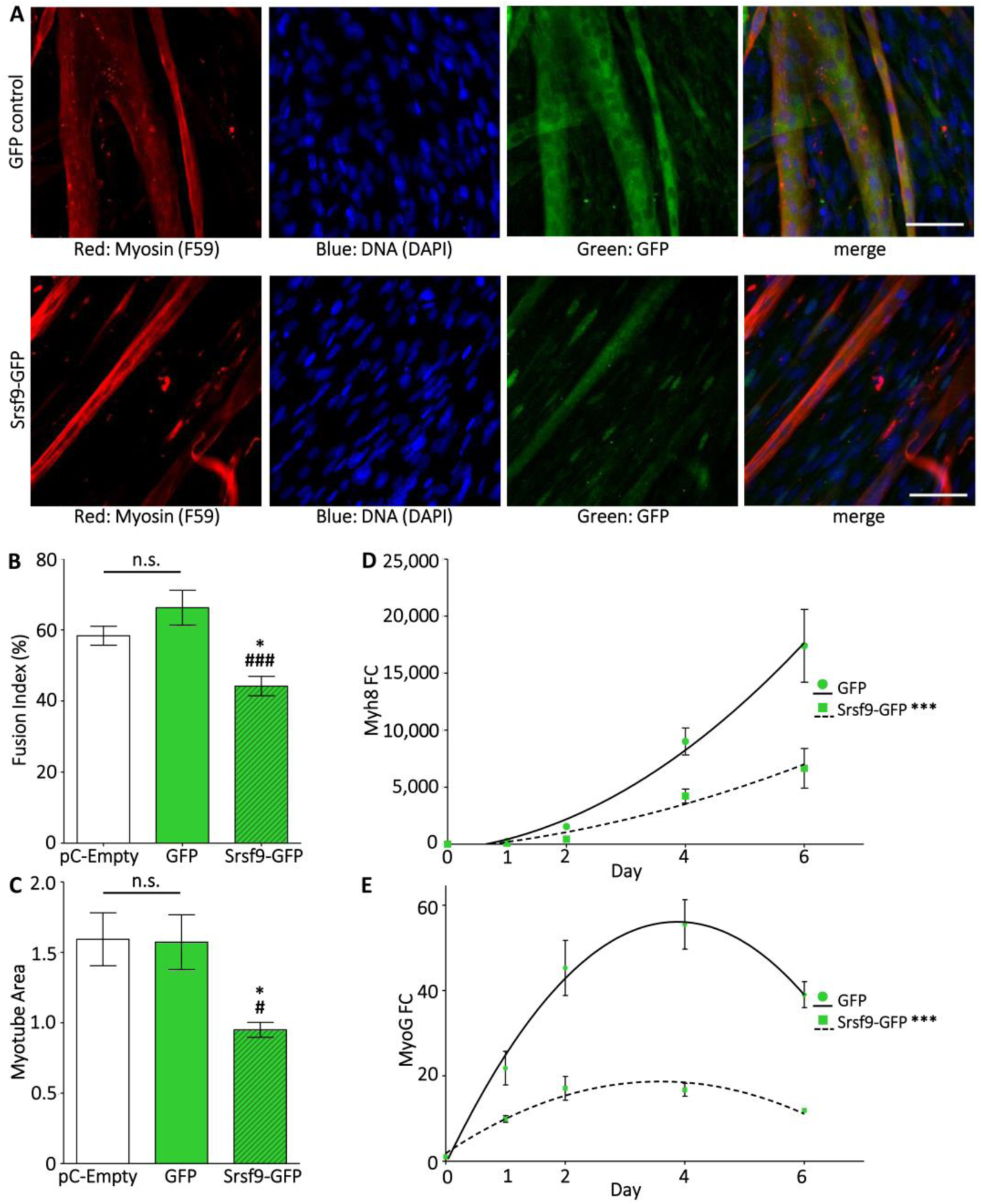
mir-1/206-resistant Srsf9-GFP expression blunts myogenic differentiation. **(A)** C2C12 cell lines stably expressing GFP alone (GFP control) or GFP-tagged Srsf9 linked to a heterologous miR-1/206-resistant 3’ UTR (Srsf9-GFP) were differentiated for 6 days, immunostained, and imaged (20X objective). Myosin heavy chain, a marker of terminal differentiation, was visualized with the F59 antibody and an AlexaFluor568-linked secondary antibody. DNA was visualized by DAPI staining and GFP or Srsf9-GFP was visualized by GFP autofluorescence. Scale bars denote 6.5 μm. **(B)** Srsf9-GFP persistence reduces fusion. The fusion index of Srsf9-GFP cells was significantly lower than either pC-Empty cells (an empty expression plasmid) or GFP control cells. 8-10 non-overlapping fields of view were analyzed for each cell line. * = p < 0.05 vs pC-Empty; ### = p < 0.001 vs GFP control. **(C)** Srsf9-GFP persistence reduces myotube size. Myotube area was assessed by calculating the myosin-positive area of D6 myotubes. The myosin-positive area/DAPI-positive area was calculated for 8-10 non-overlapping fields of view. Myosin positive area was significantly lower than both pC-Empty and GFP control cell lines. * = p ≤ 0.05 vs pC-Empty; # = p < 0.05 vs GFP control **(D)** Srsf9-GFP persistence reduces myosin mRNA expression. mRNA levels of perinatal myosin (Myh8) were assessed by qPCR at Days 0, 1, 2, 4, and 6 of differentiation. Statistically different quadratic curves were fit to GFP control and Srsf9-GFP series. **(E)** Srsf9-GFP persistence reduces MyoG induction. MyoG mRNA levels were measured by qPCR as in Panel D. Statistically different quadratic curves were fit to GFP control and Srsf9-GFP. For D and E: asterisks next to the Srsf9-GFP legend denote p < 0.0001 for the curve fit comparison.

## Discussion

In this report we used a computational analysis of differentiating C2C12 myoblasts to identify miR-1/206 targets relevant to myogenesis. We found that the 3’ UTRs of several RNA-binding proteins, which belong to the SR splicing factor family, were highly enriched in our bioinformatics assessment. Accordingly, we discovered that down-regulation of one of its members, Srsf9, controls myoblast differentiation. One limitation of our computational method is that it predicts miR-1/206 targets based on the decrease of their RNA levels; targets that are translationally inhibited but not regulated at the mRNA level, such as Hdac4 and Igf1 [25,45,46], escaped our analysis. However, several genome-wide studies on miRNAs, which include miR-1, have shown good correlation between RNA and protein levels [28,38].

miR-1 and miR-206 are highly conserved members of the myomiR family that also includes miR-133a/b, miR-208a/b, miR-486, and miR-499. This ensemble of miRNAs, specifically expressed in cardiac and skeletal muscle, governs muscle differentiation, maintenance, and plasticity [47]. To date, several reports have shown that miR-1/206 control the expression levels of genes implicated in transcription (Pax3, Pax7, Hdac4; [24,40,45,48,49]) and cellular proliferation (Pola1; [26]). In this study, we found that down-regulation of the splicing factor Srsf9 by miR-1/206 targeting is also essential for proper C2C12 differentiation. Srsf9 cooperates with several other RNA binding proteins to repress or activate regulated splicing. It modulates the inclusion of SMN exon 7 [50], the alternative splicing of glucocorticoid receptor beta and gonadotropin-releasing hormone [51,52], inclusion/exclusion of tau exons 2 and 10 [53,54], and generation of the hnRNP A1^B^ isoform [55,56]. Notably, the main hnRNP A1 isoform regulates the alternative splicing of pyruvate kinase-M (PK-M). While the PK-M1 isoform is highly enriched in differentiated myotubes, myoblasts express the PK-M2 isoform which gives them a proliferative advantage [57]. Thus, it is tempting to speculate that, by controlling the shift in pyruvate kinase isoform expression, the combined activity of Srsf9 and miR-1/206 during differentiation lays the groundwork for metabolic adaptation. Moreover, it has been proposed that Srsf9 can interact cooperatively or compete with other splicing regulators for binding to high-affinity sites present in alternatively spliced pre-mRNAs [56]. Therefore, even a small decrease in Srsf9 cellular concentration could lead to profound changes in global splicing patterns. The SR protein cellular function is still not completely defined. Global mapping of RNA targets determined by uv crosslinking and immunoprecipitation (CLIP/iCLIP) has been carried out for Srsf1, 2, 3, and 4, and the eight drosophila SR homologs [36,58,59] but not for Srsf9. These studies performed in mouse embryo fibroblasts (MEFs) and P19 embryonal carcinoma cells revealed intriguing findings. For example, i) specific alternative splicing patterns apparently controlled by a single SR protein actually hinge upon an intricate network of relationships with many other SR proteins; ii) SR proteins can control gene expression by binding intronless transcripts as well as ncRNAs. Since no such map is available for Srsf9, it is difficult to predict through which molecular pathways Srsf9 exerts control over myoblast differentiation. Srsf9 high-affinity binding sites have been identified from a randomized pool of RNA sequences by a SELEX approach [60]. However, a computational survey of SELEX-derived consensus sequences, which are short and degenerate, revealed frequent over-representation in RNAs [61].

miR-1 and miR-206 have both been proposed as tumor suppressors. Interestingly, Srsf9 depletion reduces viability of prostate cancer cells [62]. Moreover, in two human bladder cancer cell lines, Srsf9 down-regulation by miR-1 overexpression resulted in significant reduction in cell proliferation, migration, and invasion [63]. Accordingly, siRNA-mediated Srsf9 knockdown promoted apoptosis [63,64]. These data strongly suggest that in bladder cancer cells, the tumor suppressive activity of miR-1 triggers apoptosis through direct Srsf9 inhibition. However, we did not observe any impact on cell survival when Srsf9 expression was stabilized during C2C12 differentiation.

## Conclusions

We report here that miR-1/206 target the Srsf9 3’ UTR during C2C12 myoblast differentiation. This is the first report showing a direct correlation between this member of the SR protein family and muscle maturation. We demonstrate that Srsf9 down-regulation is necessary to achieve robust cell fusion and muscle-specific gene expression. Based on the complex web of functional interactions occurring amongst the RNA splicing proteins, we suggest that the persistence of Srsf9 alters several alternative splicing events and impairs proper production of specific protein isoforms driving myoblast maturation. The data presented also emphasize the importance of maintaining appropriate miRNA-mediated gene repression in adult muscle to avoid re-expression of fetal genes that could cause muscle disease.

## List of abbreviations

DAPI: 4’,6-diamidino-2-phenylindole;
DAVID: Database for Annotation, Visualization, and Integrated Discovery;
dpc: days post coitum;
FC: fold change;
GFP: green fluorescent protein;
MRE: miRNA response element;
PBS: phosphate-buffered saline;
qPCR: quantitative PCR;
RNA-Seq: RNA sequencing;
SR protein: serine/arginine-rich protein;
TBS: Tris-buffered saline;
UTR: untranslated region;

## Declarations

### Ethics approval

All animal work was approved by the University of Colorado Boulder Institutional Animal Care and Use Committee under protocols 1002.07 and 1002.08

### Availability of data

All data generated or analyzed during this study are included in this published article and its supplementary files.

### Authors’ contributions

KKB and LAL conceived and designed the study. KKB performed bioinformatics analyses, luciferase assays, and stable cell line generation and accompanying analysis, analyzed data, and co-wrote the manuscript. MB was a major contributor to data analysis and co-wrote the manuscript. EKP performed mutagenesis on reporter gene constructs and performed luciferase assays. MMP assisted with cloning of reporter gene constructs. All authors read and approved the final manuscript.

## Acknowledgments

We thank Drs. John and Christine Seidman for technical assistance with RNA-Seq data collection and analysis. We thank Dr. Joe Dragavon and the BioFrontiers Advanced Light Microscopy Core facility for technical support during image collection. We thank Dr. Eunhee Chung for assistance with embryonic tissue collection.

## Supplementary Figures

**Supplementary Figure 1.** Supplementary Figure 1.tif

**Representative masked images used for D6 myotube area calculation. (A)** Myosin heavy chain, a marker of terminal differentiation, was visualized with the F59 antibody and an AlexaFluor568-linked secondary antibody. **(B)** DNA was visualized by DAPI staining. **(C)** The binary myosin mask corresponding to Panel A. **(D)** The binary DNA mask corresponding to Panel B.

**Supplementary Figure 2.** Supplementary Figure 2.tif

**miR-1 and miR-206 levels increase during C2C12 differentiation.** Expression was assessed by qPCR and normalized to sno202 in growing myoblasts (D0) and days 1, 2, 4, and 6 of differentiation. Fold changes vs D0 are presented. * = p < 0.05; *** = p < 0.001

**Supplementary Figure 3.** Supplemental Figure 3.tif

**Non-myogenic miR-450a-1 is processed and active when ectopically expressed in myoblasts.** C2C12 myoblasts were co-transfected with a reporter construct containing two tandem copies of the reverse complement of miR-450a-1 (2×450a-1) in the 3’ UTR along with expression constructs for miR-206, miR-1, or miR-450a-1. Cells were harvested 24 hours later. Firefly-normalized Renilla luciferase activity was equivalent between miR-206 and miR-1 expression but significantly down-regulated in the presence of miR-450a-1. *** = p < 0.001 vs miR-206; ### = p < 0.001 vs miR-1 N = 3 independent cultures for each.

**Supplementary Figure 4.** Supplementary Figure 4.tif

**miR-1/206-resistant Srsf9-GFP expression does not change during differentiation of stable cell lines. (A)** GFP or Srsf9-GFP mRNA levels in the corresponding stable C2C12 cell lines were compared to a negative control C2C12 cell line (pC-Empty) incorporating the empty expression vector pCDNA3.1(-). Expression was assessed by qPCR with GFP-specific primers and normalized to 18S rRNA levels. Fold change relative to pC-Empty is presented. N = 3 independent cultures for each. **(B)** Srsf9-GFP protein is expressed in the stable cell line. Expression in pC-Empty, GFP, and Srsf9-GFP cells was assessed by western blot with a GFP-specific antibody. **(C)** Srsf9-GFP mRNA levels are stable during differentiation of the Srsf9-GFP cell line. Expression was assessed by qPCR as in A. There is no statistical difference amongst the time points.

**Supplementary Table 1.** Supplementary Table 1.xlsx

**Primer sequences.** All primer sequences are presented 5’ → 3’.

**Supplementary Table 2.** Supplementary Table 2.xlsx

**C2C12 RNA–Seq data.** mRNA from C2C12 day 6 myotubes (AD1, AD2, AD3) and proliferating myoblasts (BD1, BD2, BD3) was sequenced with a paired end protocol on an Illumina HiSEQ. The table presents reads per million aligned to the mouse mm9 genome. Column A is the UCSC Gene Symbol and Columns B-E list chromosomal coordinates and coding strand designation.

**Supplementary Table 3.** Supplementary Table 3.xlsx

**TargetScan-predicted miR-1/206 targets that decrease during C2C12 differentiation.** Predicted mouse miR-1/206 targets from TargetScan v. 7.2 were crossed with all mRNAs that decreased by 1.3-fold or greater during C2C12 differentiation. This filtered the list of all 896 candidate targets to 354 that were down-regulated at the mRNA level. This is the list that was used for functional clustering gene ontology analysis with the DAVID database. Fold change from D0 to D6 is presented along with MRE scores from TargetScan.

**Supplementary Table 4.** Supplementary Table 4.xlsx

**Functional Clustering output for DAVID analysis of 354 candidate miR-1/206 targets.** Candidate miR-1/206 targets were uploaded to DAVID and the functional clustering tool was applied to the gene set. 21 GO term clusters of related gene sets had an enrichment score greater than 1.3, corresponding to an average p-value of less than 0.05 for all GO categories in a cluster. Clusters are ranked by descending Enrichment Score and include all output columns from DAVID.

