## Supplementary figures and images for "miR-1/206 down-regulates splicing factor Srsf9 to promote myogenesis"

### Supplementary Figure 1

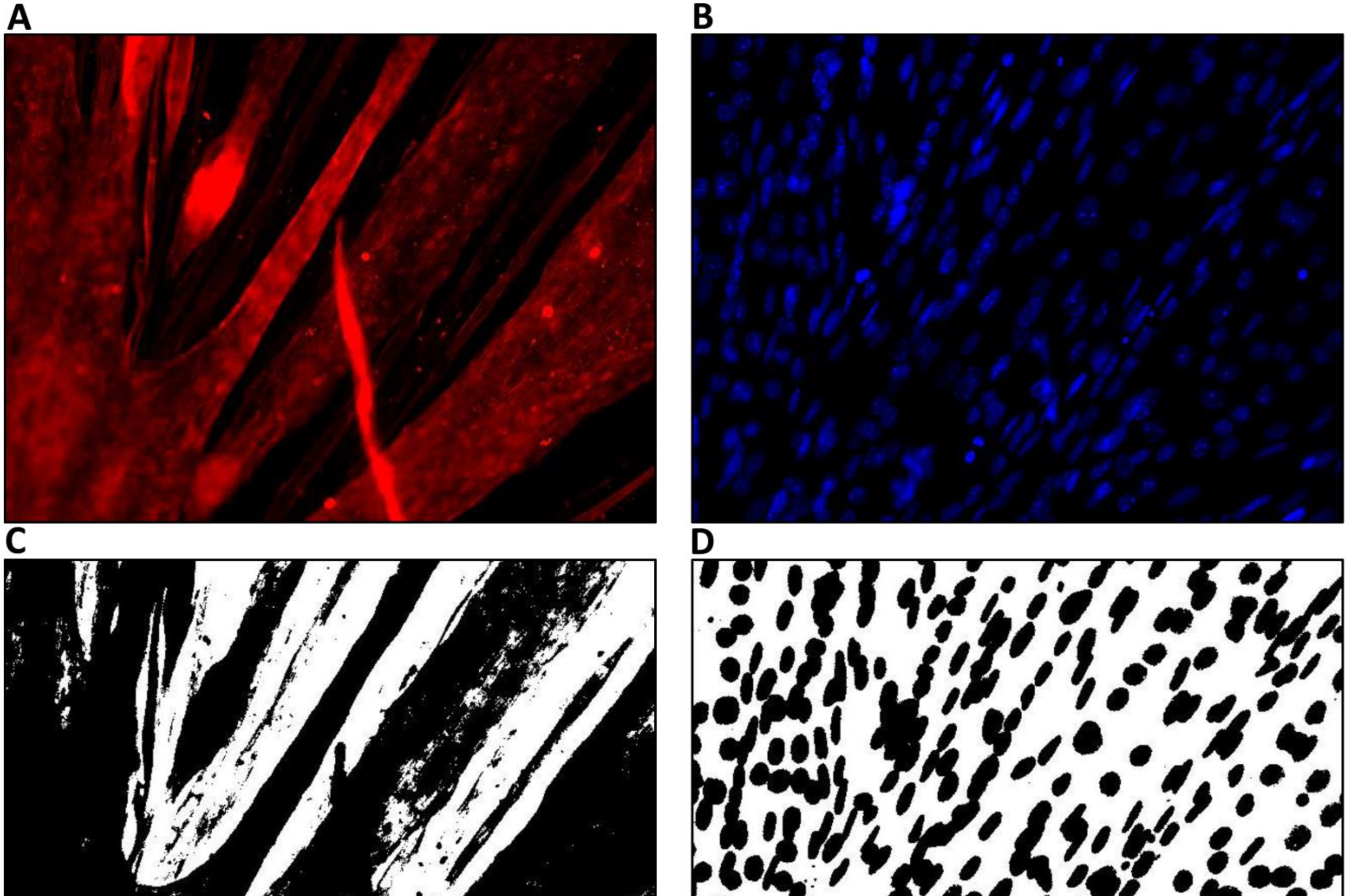

### Supplementary Figure 2

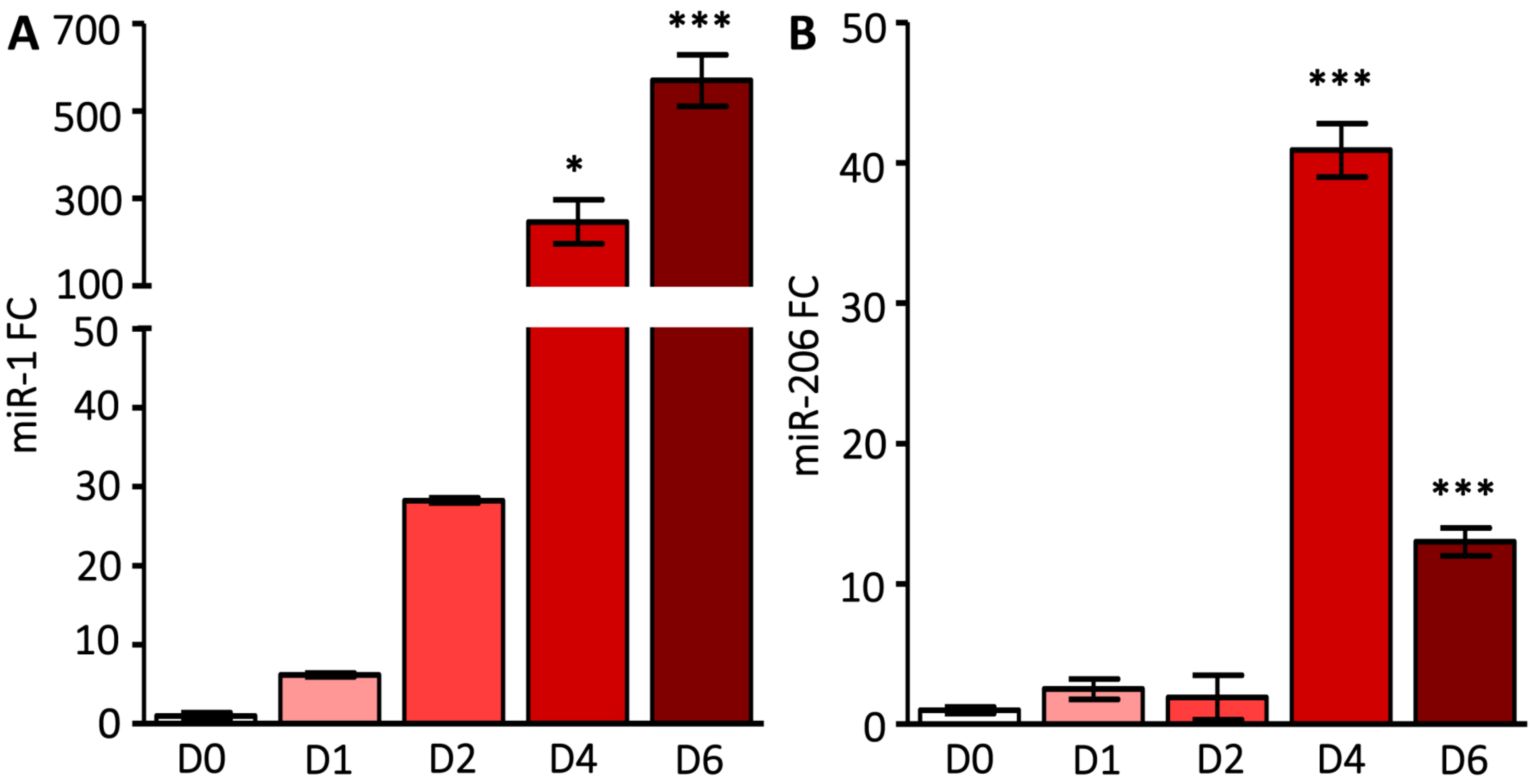

### Supplementary Figure 3

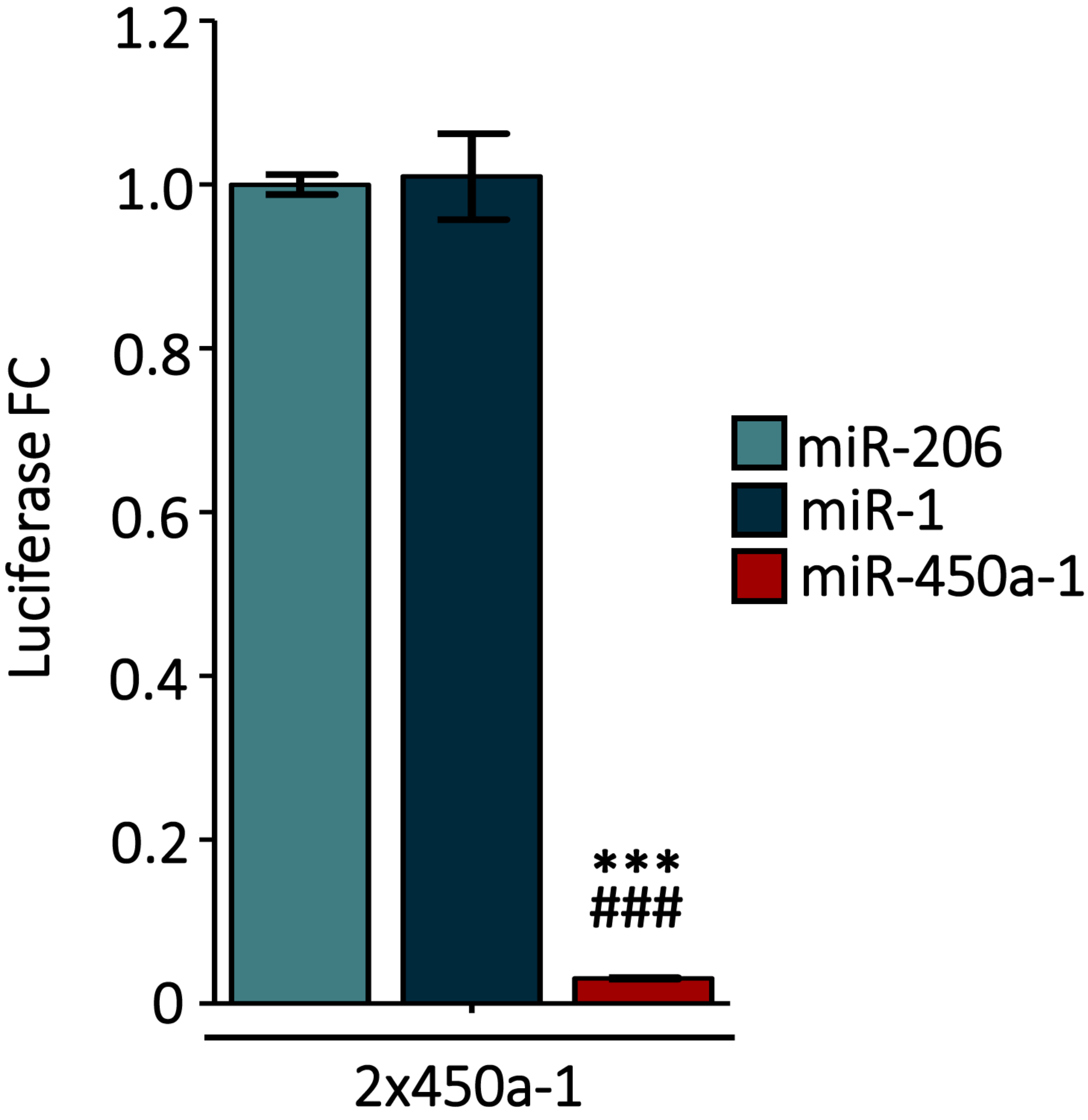

### Supplementary Figure 4

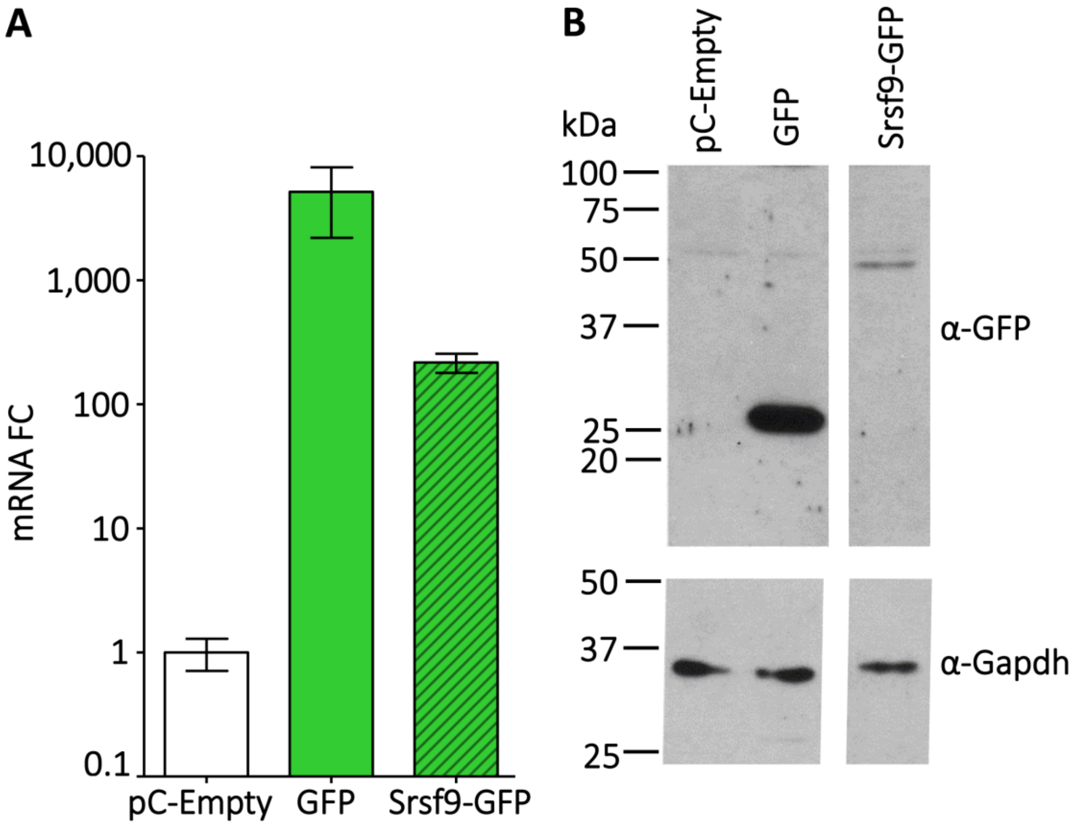
